# Distinct transcriptional programs of SOX2 in different types of small cell lung cancers

**DOI:** 10.1101/857144

**Authors:** Yuki Tenjin, Kumi Matsuura, Shinji Kudoh, Shingo Usuki, Tatsuya Yamada, Akira Matsuo, Younosuke Sato, Haruki Saito, Kosuke Fujino, Joeji Wakimoto, Takaya Ichimura, Hirotsugu Kohrogi, Takuro Sakagami, Hitoshi Niwa, Takaaki Ito

## Abstract

*SOX2* is an oncogene in human small cell lung cancer (SCLC), an aggressive neuroendocrine (NE) tumor. However, the roles of SOX2 in SCLC remain unclear, and strategies to selectively target SOX2 in SCLC cells have not yet been established. We herein demonstrated that SOX2 is involved in NE differentiation and tumorigenesis in cooperation with ASCL1, a lineage-specific transcriptional factor, in the classical subtype of SCLC cell lines. ASCL1 recruits SOX2, which promotes INSM1 expression. Precursor SCLC lesions were established in *Trp53* (-/-); *CCSP*rtTA; tetO*Cre*; floxed*Rb1*; floxed*Hes1* mice, and the NE neoplasms induced were positive for Ascl1, Sox2, and Insm1. In contrast to the ASCL1-SOX2 signaling axis to control the SCLC phenotype in classical subtype SCLC, SOX2 targeted distinct genes, such as those related to the Hippo pathway, in ASCL1-negative, variant subtype SCLC. The present results support the importance of the ASCL1-SOX2 axis as a main subtype of SCLC, and suggest the therapeutic potential of targeting the ASCL1-SOX2 signaling axis and the clinical utility of SOX2 as a biological marker in the classical subtype of SCLC.

## Introduction

Lung cancer is the leading cause of cancer-related mortality worldwide. Small cell lung cancer (SCLC) accounts for approximately 14% of all lung cancers and is genetically considered to be one of the most aggressive malignant neuroendocrine (NE) tumors (Byers *et al*., 2015). Despite high response rates to first-line treatment, SCLC cases show rapid growth and metastasis and acquire multidrug resistance. The median survival of patients with SCLC is 7 months, and this has not markedly changed in the last few decades (Wang *et al*., 2017). The development of novel target molecules in therapies for SCLC remains limited. The findings of basic studies on the molecular mechanisms underlying small cell carcinogenesis have yet to be clarified, and advances in novel therapeutic development are expected (Gazdar *et al*., 2017).

The World Health Organization (WHO) Classification recognizes SCLC as a relatively homogeneous tumor, with pure SCLC and combined SCLC subtypes (Travis *et al.,* 2015). Approximately 30 years ago, Gazdar *et al*. reported the different forms of the “classic” and “variant” subtypes of SCLC. Classical SCLC cells are characterized by floating cell growth and distinct NE differentiation, and variant SCLC cells by adhesive growth and poor NE differentiation (Gazdar *et al.,* 1985; Carney *et al.,* 1985). Classical cell lines belong to NE high SCLC, which may be associated with the increased expression of Achaete-Scute complex homologue 1 (ASCL1), a member of the basic helix-loop-helix (bHLH) family of transcription factors. On the other hand, the variant cell lines belong to NE low SCLC, which is associated with the activation of *NOTCH*, *Hippo*, and *RE-1 silencing transcription factor* (*REST*) genes and prominent epithelial-to-mesenchymal (EMT) transition resulting in a mesenchymal phenotype (Lim JS *et al.,* 2017, Zhang *et al.,* 2018). ASCL1 was previously shown to be expressed at a high frequency in SCLC (Ball *et al.,* 1993). Furthermore, the knockdown of *ASCL1* induced growth inhibition and apoptosis in SCLC cell lines (Osada *et al.,* 2005, 2008). Insulinoma-associated protein 1 (INSM1) is a crucial regulator of NE differentiation in lung cancer, and is specifically expressed in SCLC, along with ASCL1 (Fujino *et al.,* 2015). Borromeo *et al*. (2016) reported that Ascl1 played a pivotal role in tumorigenesis in mouse models of SCLC, and also suggested that *SOX2* and *INSM1* were target genes of ASCL1. In human SCLC, *SOX2* was recognized as an oncogene because *SOX2* amplification was detected in some SCLCs, and *SOX2* gene suppression inhibited the cell proliferative capacity of SCLC cell lines (Rudin *et al.,* 2012).

The major aim of the present study is to elucidate the mechanisms by which SOX2 affects the phenotype and heterogeneity of SCLC. We hypothesized that SOX2 may contribute to distinct transcriptional programs and biological characteristics in both the classical and variant subtypes of SCLC. To test this hypothesis, the present study was designed to investigate the following: (1) a comparison of the target genes of SOX2 in human SCLC cell lines by a chromatin immunoprecipitation sequence (ChIP-seq) analysis. (2) the relationship between ASCL1 and SOX2 in SCLC cell lines, surgically-resected tissues, and mouse SCLC precursor lesions, and (3) the functional difference in SOX2 between the classical and variant subtypes of human SCLC cell lines. We herein demonstrated that SOX2 was more strongly expressed in some SCLC than non-SCLC (NSCLC). SOX2 regulates distinct transcriptional programs in both the classical and variant subtypes of SCLC, and, in the classical subtype, functions in an ASCL1-dependent manner.

## Results

### SOX2 is overexpressed and targeted distinct genes between classical and variant subtypes of SCLC cell lines

To examine SOX2 expression patterns, we performed a Western blot (WB) analysis of 12 lung cancer cell lines (7 SCLCs, 3 adenocarcinomas (ADCs), and 2 squamous cell carcinomas (SCCs)). The WB analysis revealed that the SOX2 protein was generally expressed at markedly higher levels in all SCLC cell lines examined than in the NSCLC cell lines. This result suggested that SOX2 plays a more pivotal role in SCLC cell lines. Four out of the 7 SCLC cell lines simultaneously expressed SOX2, ASCL1, and INSM1 (Fig. 1A). There are two subtypes of SCLC cell lines, classical and variant, which are distinguished by their morphologies or NE properties. The loss of the expression of a master transcriptional factor, such as ASCL1, is often associated with the properties of variant SCLC cell lines (Zhang *et al.,* 2018). In the present study, H69, H889, SBC1, and H1688 belonged to the classical subtype of SCLC cell lines, which were positive for ASCL1 and INSM1. In contrast, H69AR, SBC3, and SBC5 were classified as the variant subtype of SCLC cell lines, which were negative for ASCL1. We then conducted ChIP-seq to analyze SOX2 target genes in the H69, H889, and SBC3 cell lines. We also combined the results of the RNA-seq analysis for these cell lines, which showed the global expression levels of mRNAs. As shown in a Venn plot, we identified 346 SOX2-bound genes (shared in NCI-H69 and NCI-H889) that were specifically occupied in the classical subtypes and expressed at higher levels in the classical subtypes than in the variant subtype (Fig. 1B). Some neuron-related genes, such as *INSM1*, *SEZ6L*, or *SV2B*, were included as their common target genes. On the other hand, we identified 825 SOX2-bound genes (targeted in SBC3) that were specifically occupied in the variant subtype and expressed at higher levels in the variant subtype than in the classical subtypes. Hippo pathway-related genes, such as *YAP1, WWTR1*, *LATS2,* and *REST*, which were not contained in the classical subtypes, were identified in this category (Fig. 1B). To validate functional differences in SOX2 between the classical and variant subtypes of SCLC, we used the Database for Annotation, Visualization and Integrated Discovery (DAVID) online bioinformatics tool for a GO functional analysis and extracted the top 10 enriched categories in biological processes. Neuron-related categories were more likely to be included in the classical subtype, and cell development- or movement process-related categories in the variant subtype (Fig. 1C). These results suggest that SOX2 regulates distinct transcriptional programs between the classical and variant subtypes of SCLC cell lines.

**Fig. 1:**
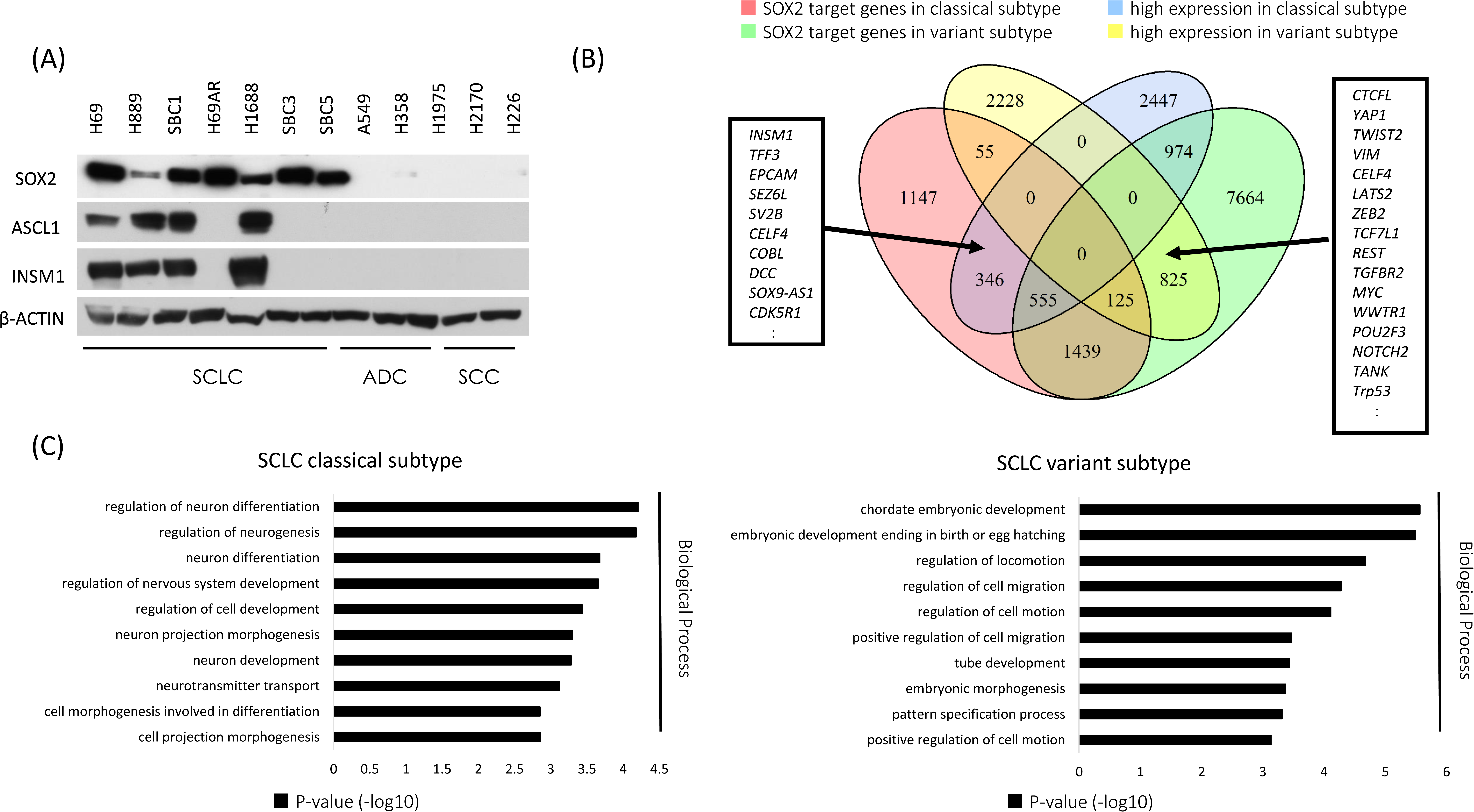
(A) WB analysis of SOX2, ASCL1, and INSM1 in lung cancer cell lines, including small cell lung carcinoma (SCLC), adenocarcinoma (ADC), and squamous cell carcinoma (SCC). SOX2 was more strongly expressed in all SCLC cell lines than in the NSCLCs examined. Four out of the 7 SCLC cell lines simultaneously expressed ASCL1 and INSM1. β-ACTIN served as an internal control. (B) ChIP-seq and RNA-seq were conducted to analyze putative SOX2 target genes and mRNA expression in the H69, H889, and SBC3 cell lines. As shown in a Venn plot, 346 specific SOX2-bound genes, which also significantly expressed mRNAs in the classical subtype, were identified (shared in NCI-H69 and NCI-H889). *INSM1* was included as a common gene. In a similar manner, 825 specific SOX2-bound genes that also significantly expressed mRNAs were identified in the variant subtype of the SBC3 cell line. Hippo pathway-related genes and *REST* were included as common genes. (C) Results of the GO functional analysis. The top 10 enriched categories in biological processes in each of the classical and variant subgroups were shown. The distinct functions of SOX2 in each subgroup were suggested.

### ASCL1 is one of the driver molecules of SOX2 and recruits SOX2 for distinct transcriptional regulation in classical subtypes of SCLC

To investigate the biological effects of ASCL1 on SOX2 in lung cancer cell lines, we performed *ASCL1* transfection experiments. An A549 cell line that stably expresses ASCL1 was established, as previously reported (Ito *et al.,* 2017; Tenjin *et al.,* 2019). A549 is a representative ADC cell line that does not express ASCL1 or other NE markers. A549 expressed SOX2 more weakly than the SCLC cell lines examined (Fig. 1A). We detected the up-regulated expression of SOX2 following the forced expression of ASCL1 in A549 cells. The expression of INSM1 and WNT11 was also induced (Fig. 2A). Immunohistochemical staining (IHC) revealed that SOX2 was more strongly expressed in xenotransplanted tumor cells with the *ASCL1* transgene than in those from A549 mock cells (Fig. 2B). To show changes in transcriptional regulation driven by SOX2 in an ASCL1-dependent manner, we compared the target genes of SOX2 between *ASCL1*-transfected A549 and A549 mock cells. We combined the results of ChIP- and RNA-seq analyses for these cells and identified 35 genes that had specific SOX2-binding peaks and higher expression levels of mRNAs in *ASCL1*-transfected A549 cells than in mock cells (Fig. 2C). This result suggested that SOX2 drives the distinct transcriptional regulation of SCLC. An integrated genome viewer (IGV) snapshot showed SOX2 binding at the overlap region of the transcription start site (TSS) of *INSM1* (Fig. 2D). To confirm that ASCL11 actually affects SOX2 expression in SCLCs, we conducted *ASCL1* knockdown experiments using RNA interference (RNAi) in H69, H889, and SBC1 cell lines as representatives of SCLC cells that simultaneously express ASCL1 and SOX2. The knockdown of ASCL1 expression in these cells resulted in significant reductions in SOX2 in 2 out of the 3 cell lines, namely, H889 and SBC1 (Fig.2E). Furthermore, to examine SOX2, ASCL1, and INSM1 expression patterns, we IHC stained 30 surgically resected SCLC tissues for these proteins as well as 20 surgically resected ADC and 20 surgically resected SCC tissues for SOX2 and ASCL1. IHC revealed that SCLCs and SCCs expressed SOX2 at slightly higher levels than ADCs: approximately 70.0% in SCLCs, 55.0% in SCCs, and 35.0% in ADCs. In SCLC tissues, 60% of cases (18 out of 30 cases) were doubly positive for ASCL1 and SOX2 and were also positive for INSM1. Although SOX2-positive, ASCL1-negative cases were detected (3 out of 30 cases), there were no ASCL1-positive, SOX2-negative cases (0 out of 30 cases). INSM1 was strongly expressed in SCLC (25 out of 30 cases) and all ASCL1-positive cases simultaneously expressed INSM1 (18 out of 18 cases) (Fig. 2F, Table 1, and Supplementary Fig. S1). Furthermore, based on the results showing that SOX2, ASCL1, and INSM1 were more likely to be co-expressed in SCLC, we surveyed public datasets of gene expression profiling in human SCLC samples and examined their relationships. The RNA-seq dataset using tumor samples from 79 SCLC patients confirmed the coordinated expression of *ASCL1* and *SOX2* in human SCLC tissue samples (GSE60052: ρ=0.327759, p=0.003168). In the same manner, we confirmed the coordinated expression of *ASCL1* and *INSM1* (GSE60052: ρ=0.357944, p=0.001188). Heatmap data focusing on *SOX2*, *ASCL1*, and *INSM1* was obtained using the dataset reported by Jiang *et al*. (2016) (Fig. 2G). These results support the positive regulation of SOX2 and INSM1 expression by ASCL1.

**Fig. 2:**
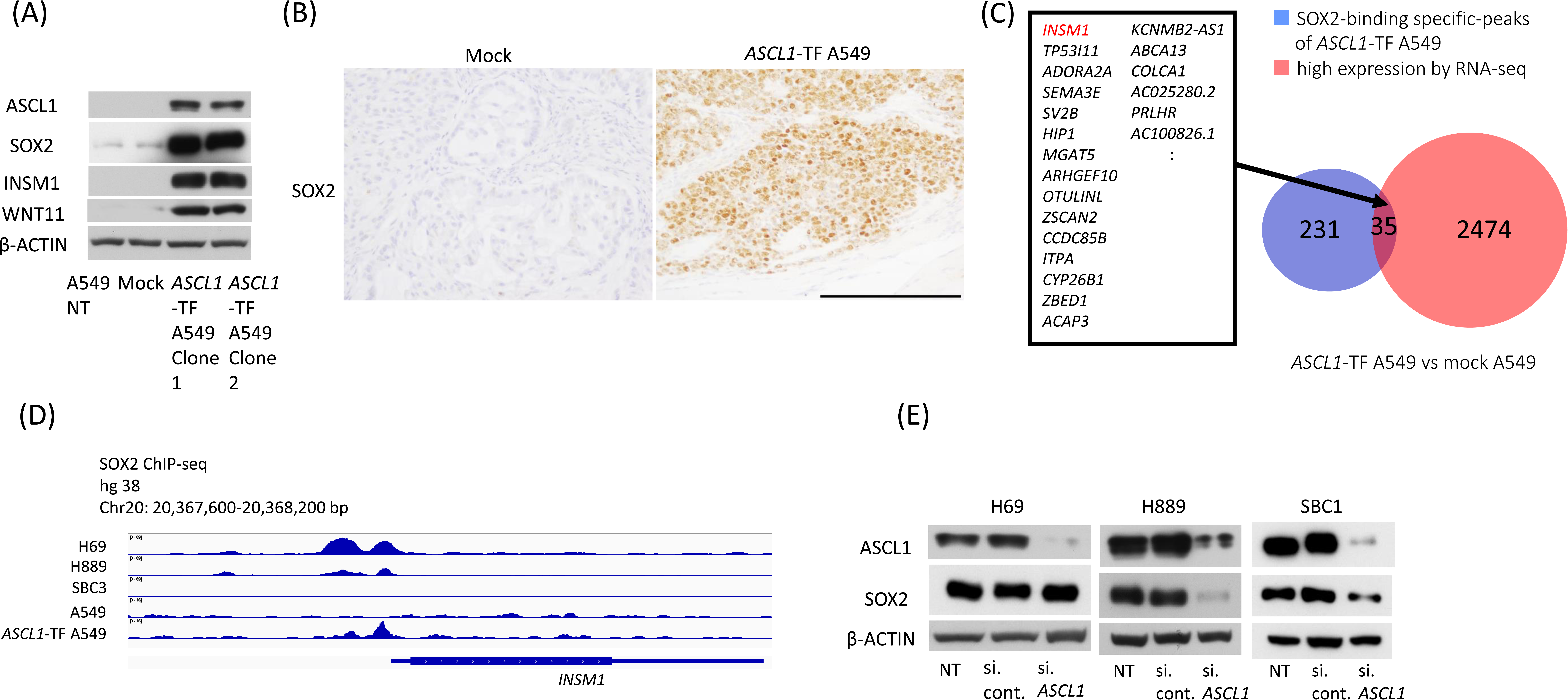
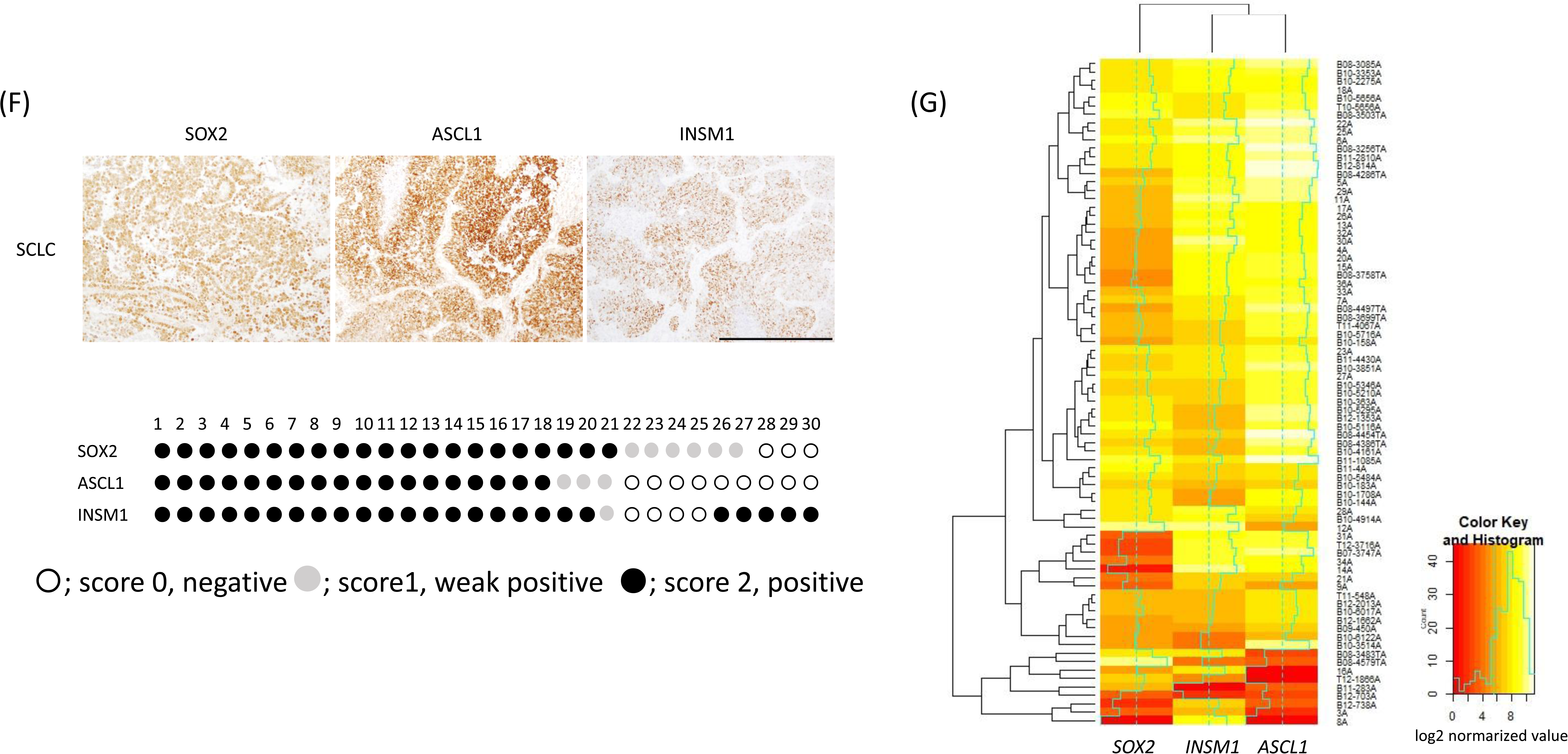
(A) A WB analysis showed that the transfection of *ASCL1* in A549 ADC cells increased SOX2, INSM1 and WNT11 expression. β-ACTIN served as an internal control. (B) Tumor tissues by the xenotransplantation of mock A549 cells and *ASCL1*-transfected A549 cells in immunodeficient mice. Using IHC, *ASCL1* transfection induced SOX2 protein expression in tumor cell nuclei. Representative images are shown. Scale bar = 200 μm. (C) Changes in transcriptional regulation driven by SOX2 between *ASCL1*-transfected A549 and A549 mock cells. ChIP- and RNA-seq combined data showed 35 specific SOX2-bound genes in *ASCL1*-transfected A549. *INSM1* was newly detected after the transfection of *ASCL1.* (D) An integrated genome viewer (IGV) snapshot showed SOX2 binding at the overlap region of the transcription starting site and ASCL1 binding near the site of the *INSM1* gene in SCLC cell lines. (E) The suppression of SOX2 by RNAi for ASCL1 was confirmed in H889 and SBC1 cells by a WB analysis (F) IHC images of surgically resected SCLC tissues for SOX2, ASCL1, and INSM1. These proteins were strongly expressed in tumor cell nuclei. Representative images are shown. Scale bar = 200 μm. (G) Expression levels of *ASCL1*, *INSM1*, and *SOX2* in the RNA-seq dataset of SCLC tissues. The GSE60052 (n=79) dataset (Jiang *et al.,* 2016) was analyzed. NT, non-treated; si, small interfering.

**Table 1.**
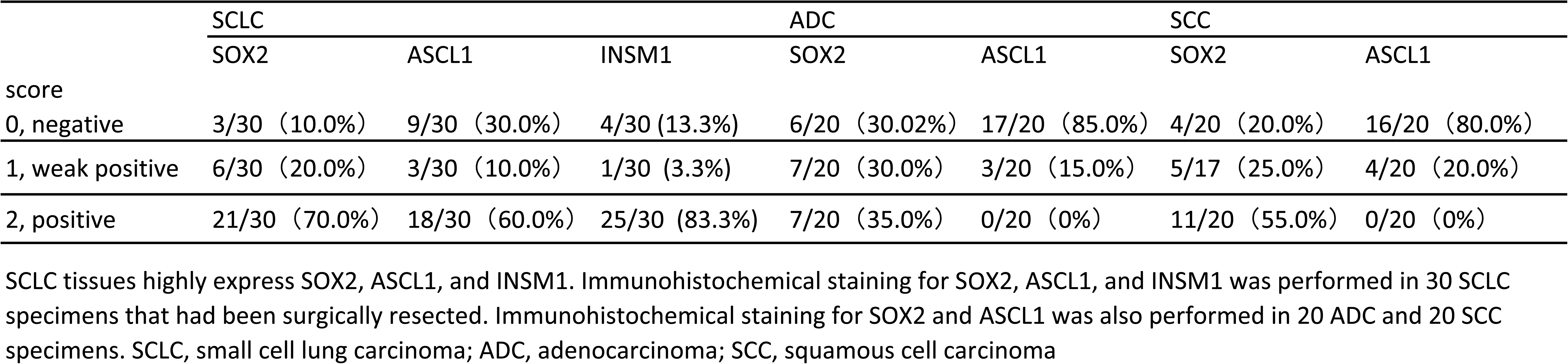
Result of Immunohistochemical Staining of Human Lung cancer

### The role of Sox2 in the classical subtype of SCLC cell lines and the *ASCL1*-transfected A549 cell line

To investigate the biological significance of SOX2 in the classical subtype of SCLC cell lines and the *ASCL1*-transfected A549 cell line, we conducted *SOX2* knockdown experiments using RNAi on these cells. The knockdown of *SOX2* expression resulted in significant reductions in *ASCL1* (Supplementary Fig. S2) and INSM1 expression (Fig. 3A) in the H69, H889, and SBC1 cell lines. Furthermore, the expression of WNT11 and CDH1 was reduced in the H69, H889, and SBC1 cell lines after the knockdown of SOX2 (Fig. 3A). This result suggests that SOX2 affected EMT in SCLC. The results of the SOX2 knockdown experiment on *ASCL1*-transfected A549 cells were also shown. Not only INSM1 and WNT11, but also NOTCH1, MYC, TCF4, RBL1, and TP53 protein levels decreased after the knockdown of SOX2. The suppression of SOX2 also affected the phosphorylation of histone H3 (p-HH3) protein levels (Fig. 3A). Cell counting assays revealed that the knockdown of *SOX2* suppressed cell growth in the H69, H889, and SBC1 cells lines (Fig. 3B). This result suggests that SOX2 positively affected cell proliferation in the classical subtype of SCLC cells and regulated the expression of key molecules for the SCLC phenotype in the presence of ASCL1.

**Fig. 3:**
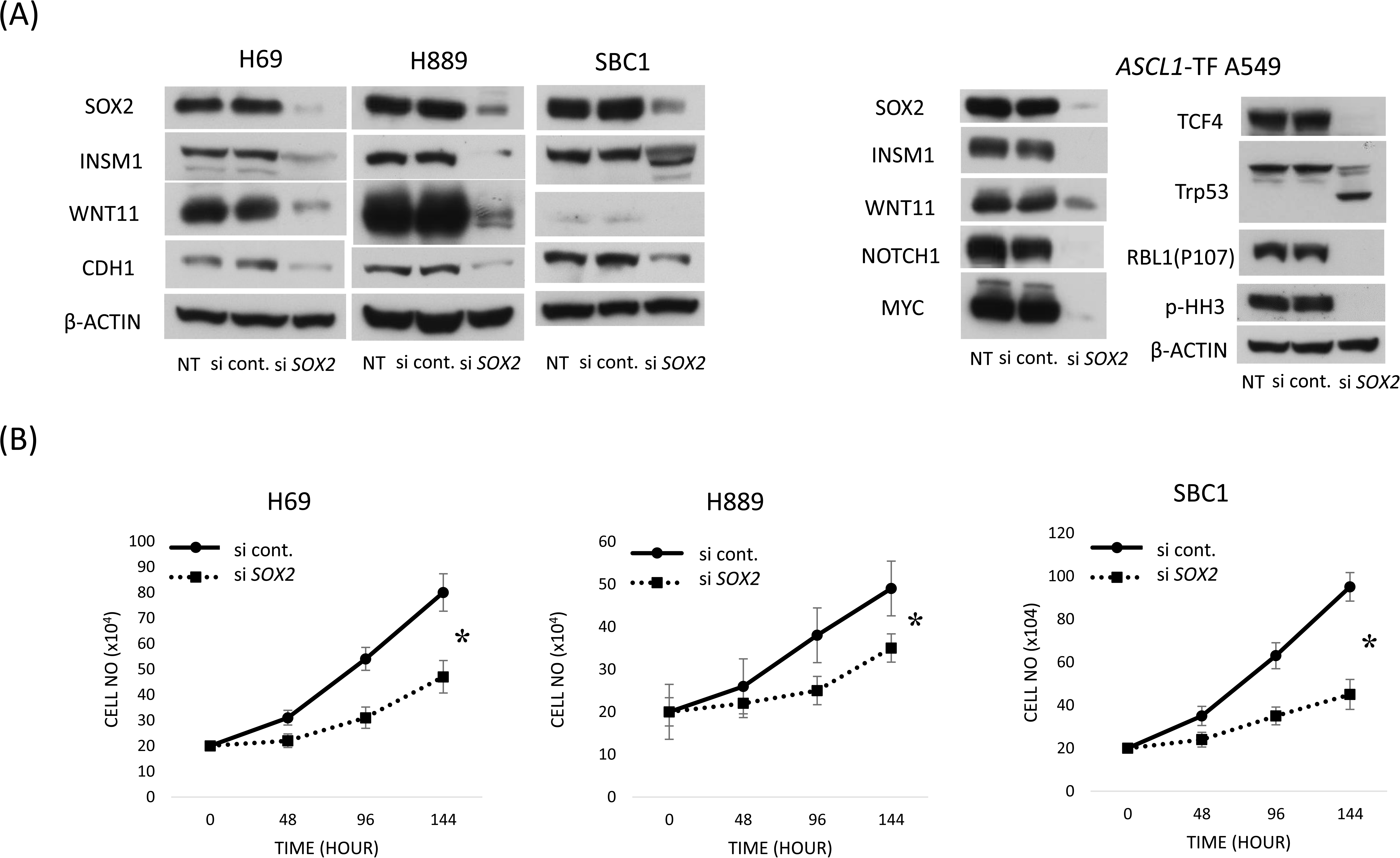
(A) SOX2 affects INSM1, WNT11, and CDH1 expression in SCLC cell lines. The suppression of INSM1, WNT11, and CDH1 was observed in the classical subtype of the SCLC cell lines, H69, H889, and SBC1s with RNAi for *SOX2*. SOX2 was also involved in the expression of NOTCH1, MYC, TCF4, Trp53, and RBL1 in *ASCL1*-TF A549 cells. (B) Cell counting assays with SCLC cell lines. The cell growth curve is shown. The suppression of cell proliferation was observed in H69, H889, and SBC1 cells with RNAi against *SOX2*. The analysis was performed in triplicate. Data are shown as the mean ±SD. Asterisks indicate a significant difference. *, p<0.05.

### The role of SOX2 in variant subtypes of SCLC cell lines

To investigate the role of SOX2 in variant subtypes of SCLC cell lines, we conducted *SOX2* knockdown experiments using RNAi on SBC3 and H69AR cell lines. SBC3 and H69AR are representative variant subtypes of the SCLC cell line that lack the expression of ASCL1 and INSM1. These cells have markedly fewer NE properties than the classical subtype of SCLC cell lines. The results obtained showed that the knockdown of *SOX2* did not significantly reduce tumor cell proliferative capacity in the SBC3 and H69AR cell lines. A quantitative real-time polymerase chain reaction (qRT-PCR) revealed that Hippo-related genes, such as *YAP1* or *TEAD1*, and *VIMENTIN* mRNA expression levels were significantly reduced after the knockdown of *SOX2*. On the other hand, the expression of *ASCL1*, *INSM1*, and *WNT11* was not significantly affected by the knockdown of *SOX2* in these cells (Fig. 4). These results suggest that SOX2 did not always sufficiently affect tumor proliferative capacity in SCLC and, in the variant subtype, it regulated the expression of downstream target genes that were distinct from those of the classical subtype of SCLC. We obtained similar results in the *SOX2* knockout experiments using the CRISPR/Cas9 system on the SBC3 and H69AR cell lines (Supplementary Fig. S3).

**Fig. 4:**
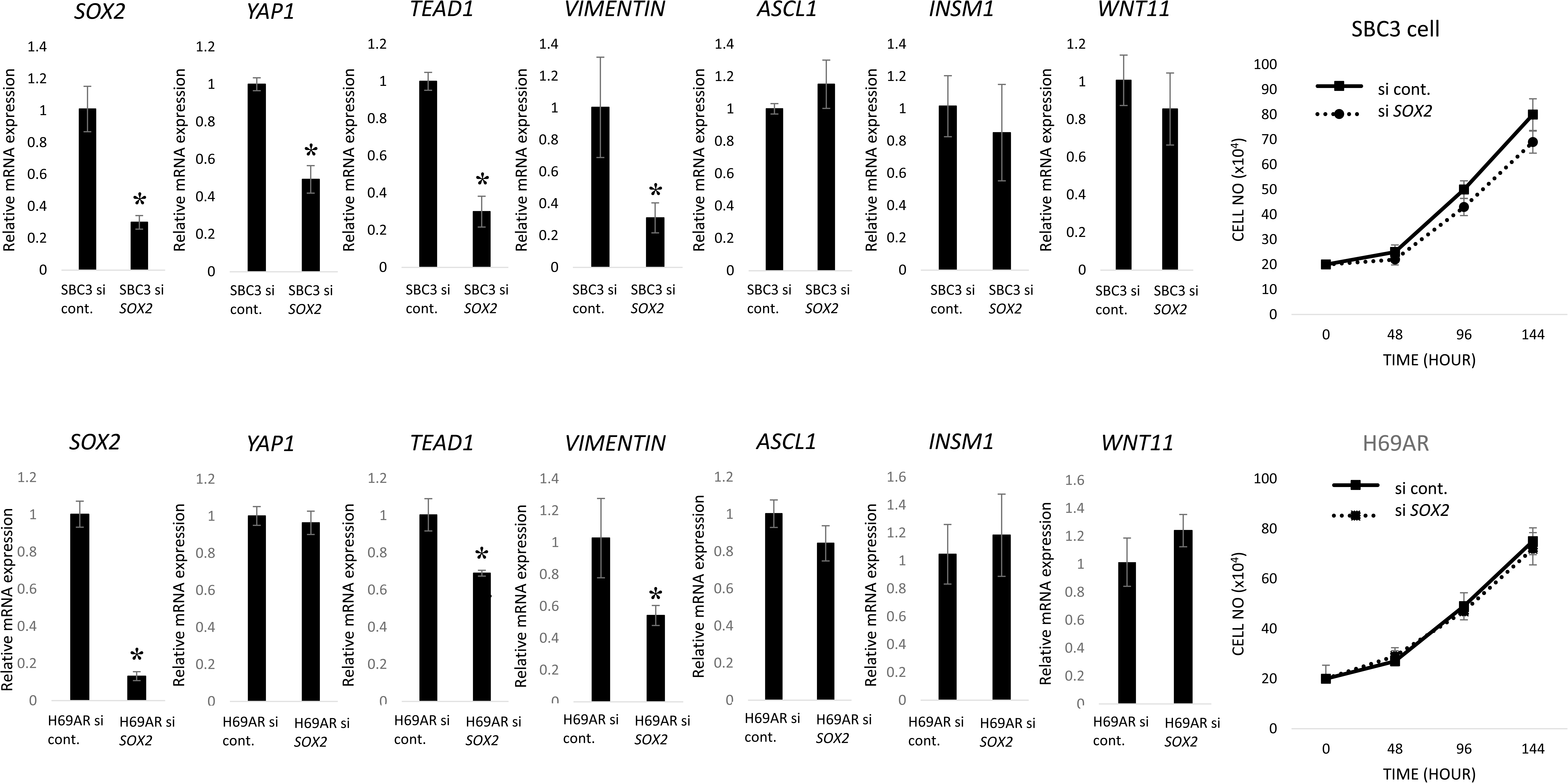
*SOX2* knockdown in the variant subtype of SCLC cell lines, SBC3 and H69AR cells, using RNAi. *SOX2* mRNA expression significantly decreased, whereas tumor proliferative capacity did not after the knockdown of *SOX2* in SBC3 and H69AR cells. *YAP1*, *TEAD1*, or *VIMENTIN* mRNA expression decreased in *SOX2*-knockout SBC3 and H69AR cells. *ASCL1*, *INSM1*, and *WNT11* mRNA expression did not change in these cell lines. The analysis was performed in triplicate. Data are shown as the mean ±SD. Asterisks indicate a significant difference. *, p<0.05.

### Ascl1, Sox2, and Insm1 were simultaneously expressed in pulmonary NE tumors in the genetically engineered mouse model

The ASCL1 highly-expressing subtype of classical SCLC represents the majority of SCLC based on multiple lines of evidence. ASCL1 is normally present in lung NE cells during development, and HES1, one of the Notch signaling targets, suppresses the expression of ASCL1 and NE cell differentiation (Borges *et al.,* 1997; Ito *et al.,* 2000). Most of the classical SCLC cell lines and primary tumor samples were shown to strongly express ASCL1 (Rudin *et al.,* 2019). Close signaling contact with the Notch-Hes1 pathway and ASCL1 expression has been proposed to exist in small cell lung carcinogenesis (George *et al.,* 2015). On the other hand, the *Trp53* and *Rb1* double-knockout mouse is fundamentally presented as a gene-engineered SCLC mouse model (Meuwissen *et al.,* 2003). In the present study, we established a *Trp53*-knockout and *Rb1* and *Hes1* double-conditional knockout mouse model, which aimed to add the effects of the inactivation of the Notch-Hes1 pathway to the fundamental SCLC mouse model. The results obtained revealed that multiple SCLC precursor lesions developed in the broncho-bronchiolar epithelium of the mouse and IHC showed that Ascl1, Sox2, and Insm1 were positively expressed in these lesions (Fig. 5). Therefore, the inactivation of Notch signaling induced precursor lesions of the classical subtype of SCLC under *Trp53* -*Rb1-* gene-deficient conditions.

**Fig. 5:**
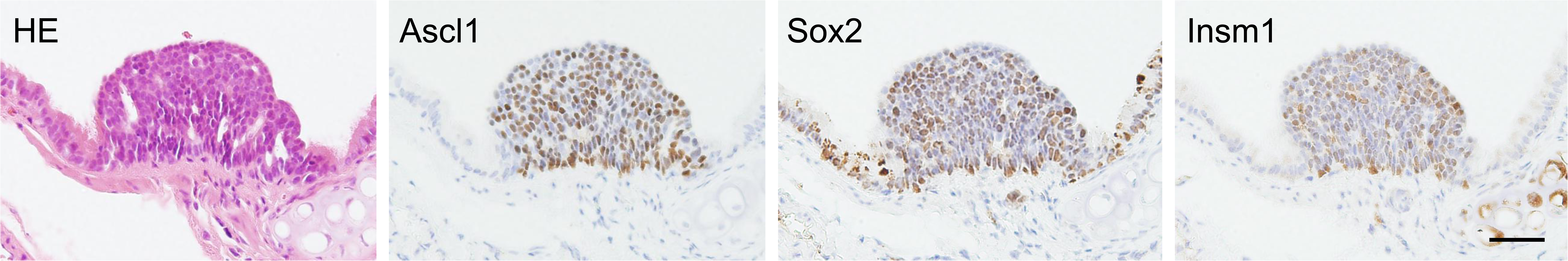
A SCLC precursor lesion in the *Trp53 (-); CCSPrtTA; tetOCre; Rb1 (fl/fl); Hes1 (fl/fl)* mouse treated with doxycycline for 6 weeks. Serial sections stained with hematoxylin and eosin (HE) and IHC for Ascl1, Sox2, and Insm1. Scale bar = 50 μm.

## Discussion

*SOX2* is an oncogene in human SCLC and its amplification has been detected in some SCLCs. In the present study, we investigated the different functions of SOX2 and whether ASCL1 was present in SCLC cell lines. SOX2 regulated INSM1 or WNT11 with the cooperation of ASCL1 in the classical subtype of SCLC. On the other hand, SOX2 targeted distinct genes, such as the Hippo pathway, in the variant subtype. These results suggest that *SOX2* is not only an oncogene in human SCLC, but may also drive distinct transcriptional regulation in each subtype of SCLC. Therefore, care is needed when considering SOX2 as a potential therapeutic target or biological marker in the diagnosis and treatment of SCLC.

We revealed that ASCL1 regulated SOX2 in the classical type of the SCLC cell line. Borromeo et al. (2016) showed that *SOX2* was one of the target genes of ASCL1 in their ChIP-seq of SCLC cell lines, which included H889 SCLC cell lines. Osada *et al*. (2005, 2008) reported roles for ASCL1 in CDH1 expression and NE differentiation. Our results support these findings, and, as a novel insight, ASCL1 and SOX2 cooperatively modulated NE differentiation or EMT in human SCLC. The knockdown of SOX2 expression in the classical subtype of the SCLC cell lines examined resulted in significant reductions in WNT11 and CDH1. In NSCLC, Bartis *et al*. (2013) reported that Wnt11 is a regulator of cadherin expression and related to the function of cellular adhesion. We previously showed that Ascl1 induced the EMT-like phenotype in A549 ADC cells (Ito *et al.,* 2017), and demonstrated that WNT11 regulated CDH1 expression in SCLC cell lines (Tenjin *et al.,* 2019). We also confirmed SOX2 binding near the site of the *WNT11* gene in the classical subtype of SCLC cell lines (data not shown). SOX2 may potently modulate NE differentiation and CDH1 expression via ASCL1 or WNT11 in SCLC.

The results obtained on the knockdown of SOX2 in *ASCL1*-transfected A549 cells are of interest. After the overexpression of ASCL1, SOX2 expression was enhanced and INSM1 and WNT11 were simultaneously expressed in this cell. The knockdown of SOX2 in *ASCL1*-transfected cells caused the suppression of INSM1 and WNT11. These results suggest that ASCL1 activates *SOX2* and regulates INSM1 and WNT11 expression together with SOX2. Furthermore, NOTCH1, TCF4, MYC, Trp53, and RBL1 expression decreased after the knockdown of *SOX2* in these cells. Notch signaling is an important cell signaling system, and the interaction between Notch receptors and their ligands induces several genes, such as *HES1*, *CCND1, MYC*, and *AKT* (Rizzo *et al.,* 2008). Intratumoral heterogeneity generated by Notch signaling has been shown to promote SCLC (Lim *et al.,* 2014). The Notch1-Hes1 pathway is a repressor of NE differentiation through the decreased expression of NE-promoting transcription factors, such as ASCL1 and INSM1 (Ito *et al.,* 2000; Ball *et al.,* 2004; Hassan *et al.,* 2014; Fujino *et al.,* 2015). Chen *et al*. (2012) reported that silencing of the *SOX2* gene reduced the tumorigenic properties of A549 cells with the attenuated expression of MYC and NOTCH1 in xenografted tumors in the NOD/SCID mouse. The canonical Wnt pathway induces MYC through TCF4 and other Wnt signal components. In addition, Wnt11 has been reported to activate both canonical and non-canonical Wnt pathways (Stewart *et al.,* 2014; Rapp *et al.,* 2017). Moreover, mouse models carrying conditional alleles for both *Trp53* and *Rb1* developed small cell carcinoma in the lung (Meuwissen *et al.,* 2003). The universal bi-allelic inactivation of *Trp53* and *RB1* was previously reported in human samples (George *et al.,* 2015). Meder *et al*. (2016) showed that NOTCH, ASCL1, Trp53, and RB alterations defined an alternative pathway driving NE and small cell carcinomas. Moreover, the ablation of all three retinoblastoma family members, Rb1, Rbl1, and Rbl2, in the mouse lung resulted in the formation of NE tumors (Lázaro *et al.,* 2017). We demonstrated the induction of precursor SCLC lesions that simultaneously expressed Ascl1, Sox2, and Insm1 in *Trp53 (-/-); CCSPrtTA; tetOCre; Rb1 (fl/fl); Hes1 (fl/fl)* mice. SOX2 has been suggested to play an important role, particularly in an ASCL1-dependent manner, in the SCLC phenotype and tumorigenesis in the interaction with these principle tumor suppressants.

We performed SOX2 knockdown or knockout experiments using RNAi or the CRISPR/Cas9 system in the variant subtype of SCLC, SBC3, and H69AR cells. After the knockdown of *SOX2, YAP1* and *TEAD1* mRNA expression levels decreased in these cells. A previous study reported that YAP1, the main Hippo pathway effector, was frequently lost in high-grade NE lung tumors, and showed reciprocal expression against INSM1 (McColl *et al.,* 2017). Among SCLC, the loss of YAP1 correlated with the expression of NE markers, and a survival analysis revealed that YAP1-negative cases were more chemo-sensitive than YAP1-positive cases (Ito *et al.,* 2016). The YAP/TAZ subgroup displayed an adherent cell morphology (Horie *et al.,* 2016) and lower expression levels of ASCL1. *REST* was also included in the genes of SOX2-bound sites in variant subtypes. REST encodes a transcriptional repressor that represses neuronal and NE genes in non-neuronal and non-NE tissues and, thus, serves as a negative regulator of neurogenesis, including SCLC (Gao *et al.,* 2011; Thiel *et al.,* 2015; Lim *et al.,* 2017). We demonstrated that SOX2 modulated the Hippo pathway in the variant subtype of SCLC in the present study. We also showed that *VIMENTIN* mRNA levels decreased after the knockdown of SOX2 in these cells. In the classical subtype of SCLC cell lines, CDH1 expression decreased after the knockdown of SOX2 by RNAi (Fig. 3A). These results suggest that SOX2 potently modulated EMT in lung cancer via the Hippo or Wnt signaling pathway in SCLC.

Rudin *et al*. (2012) reported that the knockdown of *SOX2* by doxycycline-inducible shRNA inhibited cell proliferation in SCLC cell lines. In the classical subtype SCLC cell lines, we demonstrated that SOX2-knockdown using RNAi decreased tumor cell proliferative capacity. In contrast, the knockdown of *SOX2* in SBC3 and H69AR cells did not significantly affect their cell proliferative capacity. This functional discrepancy may be attributed to functional differences in Sox2 between the classical and variant subtypes of SCLC. Furthermore, Rudin *et al*. (2019) recently proposed a nomenclature to describe SCLC subtypes according to the dominant expression of transcription factors. They divided SCLC into 4 subtypes, which considered the master regulators of SCLC; ASCL1, NEUROD1, POU2F3, or YAP1. In the present study, we revealed that SOX2 cooperated with the key regulatory molecules, such as ASCL1 and YAP1, in SCLC, and showed that SCLC may be divided into 3 groups based on the expression of ASCL1 and SOX2; ASCL1-SOX2 doubly high, ASCL1 low and SOX2 high, and ASCL1-SOX2 doubly low (Fig. 6). In addition to genomic profiling, which has been adopted in clinical practice, several research initiatives to catalog DNA, RNA, and protein profiles among lung SCC and ADC have accelerated the pace of discovery, such as The Cancer Genome Atlas (TCGA). However, similar efforts have not yet been achieved for SCLC due to the lack of adequate tumor tissue (Byers *et al.,* 2015). This needs to be investigated in a large prospective or cohort study in the future.

**Fig. 6:**
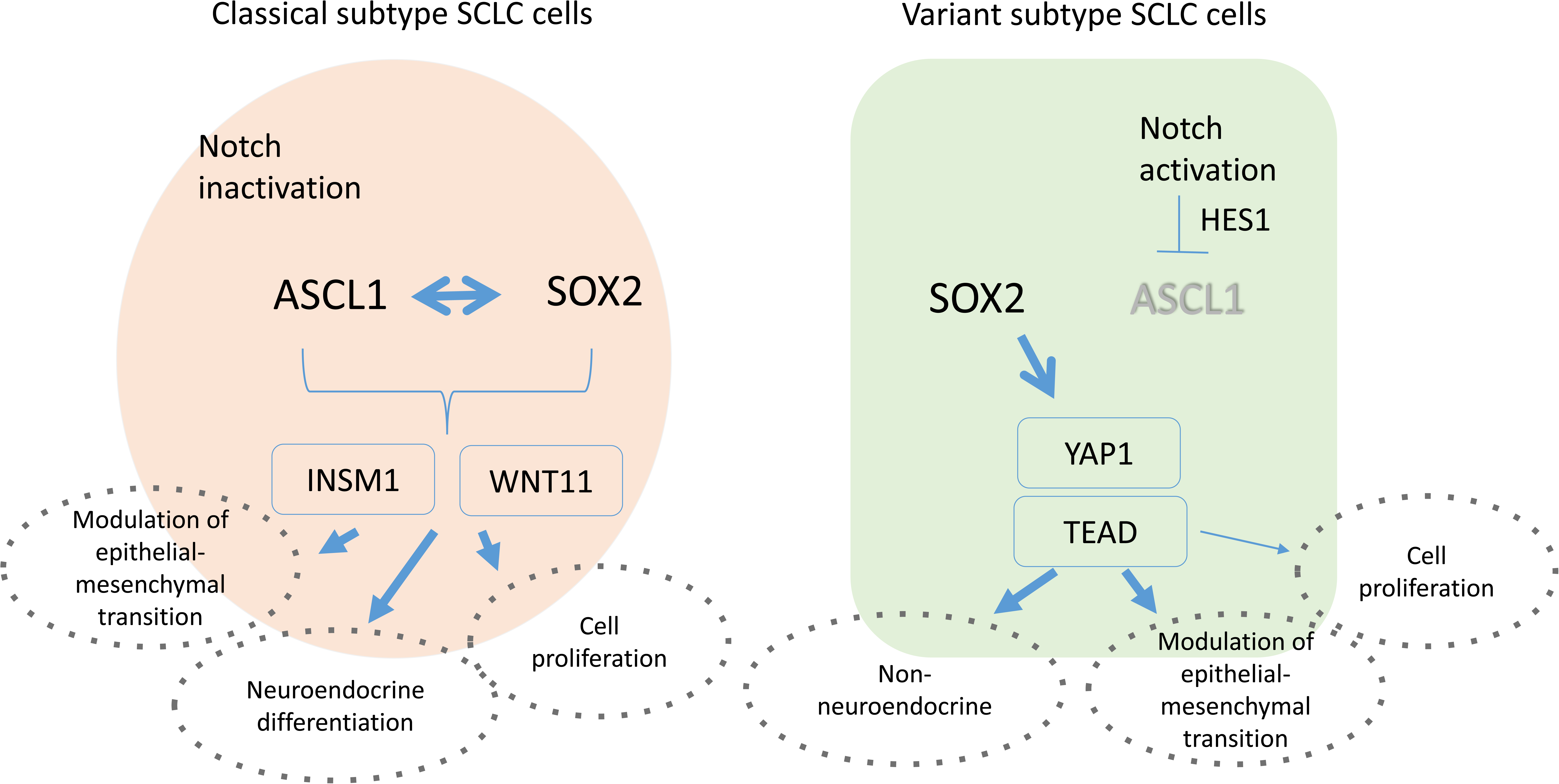
Distinct roles for SOX2 in classical and variant subtypes of SCLC were summarized. ASCL1 and SOX2 cooperatively regulate INSM1 and WNT11 expression and SOX2 affects not only tumor cell proliferation, but also EMT modulation or NE differentiation in the classical subtype. On the other hand, SOX2 affects distinct cell signaling, such as the Hippo pathway, and modulates EMT in the variant subtype. SOX2 affects tumor cell survival slightly more in the classical subtype than in the variant subtype of SCLC.

In summary, the classical subtype of SCLC frequently and strongly expresses both SOX2 and ASCL1. ASCL1-recruited SOX2 plays an important role in driving distinct transcriptional regulation. We demonstrated that SOX2 regulates lineage-specific genes, such as *INSM1*, in the classical subtype of SCLC. While we revealed the significance of SOX2 for cell growth and the modulation of EMT, we detected a functional discrepancy in SOX2 between the classical and variant subtypes of SCLC. The present results suggest that the ASCL1-SOX2 axis is extremely important as a potential therapeutic target or biological marker in the classical subtype of SCLC. On the other hand, the fundamental role of SOX2 in the variant subtype, in which ASCL1 is negative, was shown in the activation of the Hippo signaling pathway. Further studies on SOX2 that focus on highly specific molecules, for example, those that are involved in the recruitment of SOX2 in the variant subtype of SCLC, are needed. The present study promotes our understanding of the significance of SOX2 in SCLC, which will hopefully lead to the development of novel targeted therapies and better prognoses for patients with SCLC.

## Materials and Methods

### Cell Lines

Seven SCLC cell lines (H69, H889, SBC1, H69AR, H1688, SBC3, and SBC5), 3 ADC cell lines (A549, H358, and H1975), and 2 SCC cell lines (H2170 and H226) were used in the present study. H69, H889, H69AR, H1688, A549, H358, H1975, H2170, and H226 were purchased from ATCC (Manassas, VA), and SBC1, SBC3, and SBC5 from the Japan Collection of Research Bioresources Cell Bank (Osaka, Japan). All growth media were purchased from Wako Pure Chemical Industries Ltd. (Osaka, Japan) and supplied with 1% penicillin/streptomycin (Sigma–Aldrich, Ontario, Canada). A549 cells were grown in DMEM supplemented with 10% FBS (Hyclone, Logan, UT). SBC-3 cells were grown in EMEM with 10% FBS. H69, H1688, and H2170 cells were grown in RPMI 1640 medium supplemented with 2 mM L-glutamine, 10 nM HEPES, 1 mM sodium pyruvate, 4.5 g/L glucose, 1.5 g/L sodium bicarbonate, and 10% FBS. H69AR cells were grown in similar RPMI 1640 medium, but supplemented with 20% FBS. All cells were incubated at 37 °C in 5% CO_2_ and saturated humidity. Cells were maintained as subconfluent cultures before use and harvested with trypsin-EDTA (Invitrogen, Carlsbad, CA).

### Tissue Samples

We obtained tissue samples of SCLCs (n=30), ADCs (n=20), and SCCs (n=20) from the lung cancer files of the Department of Pathology and Experimental Medicine of Kumamoto University and resected at the Department of Thoracic Surgery of Kumamoto University from 70 patients for this study. A histological diagnosis of the samples was made according to the criteria of the WHO (Travis *et al.,* 2015). These sections were used for IHC staining. The present study followed the guidelines of the Ethics Committee of Kumamoto University.

### WB Analysis

Cells were prepared for a WB analysis as previously described (Yoshida *et al.,* 2013). A list of the primary antibodies used is shown in Table 2. Membranes were washed and incubated with the respective secondary antibodies conjugated with peroxidase (Amersham Pharmacia Biotech, Buckinghamshire, UK) for 1 hour, and the immune complex was visualized with the chemiluminescence substrate (Amersham Pharmacia Biotech, UK).

**Table 2.** Antibodies used for IHC and WB analysis

| Primary antibody | Manufacturer (location) | WB | IHC |
| --- | --- | --- | --- |
| Sox2 (AB5603) | Millipore (Billerica, MA) | 1:1000 | 1:100 |
| Sox2 (E-4) | Santa Cruz Biotechnology |  | 1:100<br>0 |
| ASH1 (AB15582) | Millipore (Billerica, MA) |  | 1:100 |
| MASH1 (ab213151) | abcam (Cambridge, UK) |  | 1:100<br>0 |
| Ascl1 (556604) | BD Biosciences (San Jose, CA) | 1:500 |  |
| Wnt11 (107-10576) | RayBiotech (Norcross, GA) | 1:1000 |  |
| Insm1 (C-1) | Santa Cruz Biotechnology | 1:5000 |  |
| E-cadherin (610181) | BD Biosciences | 1:1000 |  |
| Notch1 (D1E11) | Cell Signaling | 1:1000 |  |
| c-Myc (D84C12) | Cell Signaling | 1:1000 |  |
| TCF4 (D-4) | Santa Cruz Biotechnology | 1:1000 |  |
| p53 (DO-1) | Santa Cruz Biotechnology | 1:1000 |  |
| p107 (C-18) | Santa Cruz Biotechnology | 1:1000 |  |
| p-Histone H3 (Ser10) | Millipore (Billerica, MA) | 1:500 |  |
| β-actin (A-5441) | Sigma Aldrich (Oakville, ON, Canada) | 1:10000 | |

Manufacturers, quantities, and working dilutions are indicated. Ascl1, achaete-scute complex homolog-like 1; Wnt11, Wnt family member 11; Insm1, insulinoma-associated protein 1; TCF4, transcription factor 4; IHC, immunohistochemistry; WB, Western blot.

### ChIP

Three SCLC cell lines (H69, H889, and SBC3), A549 ADC cell lines, and an A549 cell line with the stable expression of ASCL1 were used in the present study. Cells were fixed in 1% formaldehyde (Thermo-Fisher) in PBS at room temperature for 10 minutes. Crosslinked cells were lysed with LB1 (50 mM HEPES-KOH (pH7.5), 140 mM NaCl, 1 mM EDTA, 10% (w/v) glycerol, 0.5% (w/v) NP-40, 0.25% (w/v) TritonX-100, proteinase inhibitor cocktail (Sigma)) and washed with LB2 (10 mM Tris-HCl (pH 8.0), 200 mM NaCl, 1 mM EDTA, 0.5 mM EGTA, proteinase inhibitor cocktail). Chromatin lysates were prepared in RIPA buffer (Thermo 89900; 25 mM Tris-HCl pH7.6, 150 mM NaCl, 0.1% SDS, 1% NP-40, 1% sodium deoxycolate, proteinase inhibitor cocktail), sonication with Covaris S220 (Peak Incident Power, 175 : Acoustic Dutu Factor, 10% : Cycle Per Burst, 200 : Treatment time, 600sec : Cycle, 6). ChIP was performed using chromatin lysates equivalent to 1.0 × 10 7 cells, and protein A Dyna-beads (Thermo-Fisher) coupled with the antibody against Sox2 (raised by us). After 4 hours of incubation at 4 °C, beads were washed 4 times in a low salt buffer (20 mM Tris-HCl (pH 8.0), 0.1% SDS, 1% (w/v) TritonX-100, 2 mM EDTA, 150 mM NaCl), and two times with a high salt buffer (20 mM Tris-HCl (pH 8.0), 0.1% SDS, 1% (w/v) TritonX-100, 2 mM EDTA, 500 mM NaCl). Chromatin complexes were eluted from the beads by agitation in elution buffer (10 mM Tris-HCl (pH 8.0), 300 mM NaCl, 5 mM EDTA, 1% SDS) and incubated overnight at 65 °C for reverse-crosslinking. Eluates were treated with RNase A and Proteinase K, and DNA was ethanol precipitated.

### ChIP -seq data analysis

ChIP-seq libraries were prepared using 20 ng of input DNA, and 1-3 ng of ChIP DNA with KAPA Library Preparation Kit (KAPA Biosystems) and NimbleGen SeqCap Adaptor Kit A or B (Roche) and sequenced by Illumina NextSeq 500 (Illumina) using Nextseq 500/550 High Output v2 Kit (Illumina) to obtain single end 75-nt reads. The resulting reads were trimmed to remove the adapter sequence and low-quality ends using Trim Galore! v0.4.3 (cutadapt v.1.15). The trimmed ChIP-seq reads were mapped to the UCSC hg38 genome assemblies using Bowtie2 v2.3.3 with default parameters. The resulting SAM files were converted to the BAM format using SAMtools v1.5. Peak calling was performed using MACS2 v2.1.1. with input DNA as a control including a q-value cut-off of 0.01 for SOX2 ChIP-seq. The distance to the nearest TSS and gene feature of the peaks were obtained from Ensembl human annotation data (GRCh38) using the annotatePeakInBatch of ChIPpeakAnno and biomaRt R packages. Peaks in the gene body were first annotated with the option ‘output=”overlapping”, and the remained peaks were then annotated to the nearest TSSs regardless of the distance between them. Protein binding sites were shown along with genomic loci from RefSeq genes on the genome browser IGV.

### RNA sequence (RNA-seq)

RNA-seq was performed by the Liaison Laboratory Research Promotion Center (LILA) (Kumamoto University) as follows. Total RNA was isolated using the RNeasy Mini Kit (Qiagen, Germany). 2100 Bioanalyzer was used to detect the concentration and purity of total RNA. All samples with an RNA integrity number (RIN) >7.5 were used for sequencing. Library DNA prepared according the Illumina Truseq protocol using Truseq Standard mRNA LT Sample Prep Kits and sequenced by Nextseq 500 (Illumina) was used for analysis, and the data were converted to Fastq files. Quality control of the data was performed by FastQC. The reads were then trimmed to remove the adapter sequence using Trim Galore! v0.5.0 (Cutadapt v 1.16), and low-quality reads were filtered out using FASTX-toolkit v0.0.14. The remaining reads were aligned to the Ensembl GRCh38.93 reference genome using STAR ver.2.6.0a. FPKM (fragments per kilobase of exon per million reads mapped) values were calculated using Cuffdiff. Significant genes were extracted by cuffdiff (p<0.05). A differential expression analysis was performed using the ExAtlas website (https://lgsun.irp.nia.nih.gov/exatlas/).

### Gene ontology (GO) analysis

GO annotation and classification were based on three categories, including biological process, molecular function, and cellular component. The Database for Annotation, Visualization, and Integrated Discovery Bioinformatics Resources 6.7 (DAVID 6.7, http://www.david.niaid.nih.gov) was used for the GO analysis (Huang *et al.,* 2009). The gene lists contained significant genes in the RNA-seq analysis and were also targeted by Sox2 in our ChIP-seq analysis. To visualize key biological processes, the DAVID online database was used. The top 10 categories in each classical and variant subgroup of SCLC were taken as excerpts for Fig. 1C.

### Transfection with siRNA

We purchased Sox2 siRNA (sc-41120) and Ascl1 siRNA (sc-37692) from Santa Cruz Biotechnology (Santa Cruz, USA) and transfected them into cells using an electroporator (NEPA21 pulse generator; Nepa Gene, Chiba, Japan) as described in the manufacturer’s instructions. These were a pool of 3 different siRNA duplexes and sequences for Sox2 were as follows. sense; 5’-GAAUGGACCUUGUAUAGAUTT −3’, antisense; 5’-AUCUAUACAAGGUCCAUUCTT −3’ (sc-38408A), sense; 5’ –GGACAGUUGCAAACGUGAATT −3’, antisense; 5’-UUCACGUUUGCAACUGUCCTT −3’ (sc-38408B), and sense; 5’-GAAUCAGUCUGCCGAGAAUTT −3’, antisense; 5’-AUUCUCGGCAGACUGAUUCTT −3’ (sc-41120C). The sequences for Ascl1 were as follows. sense; 5’-CCAACAAGAAGAUGAGUAATT-3’, antisense; 5’-UUACUCAUCUUCUUGUUGGTT-3’ (sc-37692A), sense; 5’-GAAGCGCUCAGAACAGUAUTT-3’, antisense; 5’-AUACUGUUCUGAGCGCUUCTT-3’ (sc-37692B), sense; 5’-GUUCGGGGAUAUAUUAAGATT-3’, antisense; 5’-UCUUAAUAUAUCCCCGAACTT-3’ (sc-37692B). Control siRNA (Cat# sc-36869) was used as a control. Cells were harvested 48-72 h post-transfection.

### Plasmid construction and transfection

To construct pCAG-IRES-puro-FlagHA, we replicated the *ASCL1* gene of a human *ASCL1* cDNA ORF clone and replaced it with *ASCL1*. We generated pCAG-IRES-puro-FlagHA -mock from a human *ASCL1* cDNA ORF clone by cleaving out *ASCL1*. Two plasmids were transfected into A549 cells with Lipofectamine 3000 (Invitrogen) as described in the manufacturer’s instructions. After 48 h, 1 µg/mL of puromycin (Clontech) was added to cells for 2 weeks, with a medium change every 3 days. Stably transfected resistant cell lines were cloned from each transfectant.

### Cell counting assay

A cell counting method was used to evaluate the role of Sox2 in cell proliferation. After 48 h of siRNA and control transfection, cells were stained with trypan blue and counted. H69, H889, and SBC1 cells were used and seeded at 2 × 10^5^ cells on 6-well plates. Every 2 days, cells were collected and counted, after which they were seeded into new fresh medium and left at 37 °C in 5% CO2. The counting method was continued until day 6. The criteria for cellular integrity included trypan blue exclusion, an intact nucleus, and intact cell membrane. Experiments were repeated three times separately to confirm reproducibility.

### IHC and evaluation

Formalin-fixed, paraffin-embedded specimens were cut into 4-μm-thick sections and mounted onto MAS-GP–coated slides (Matsunami Glass Ind., Osaka, Japan). After being deparaffinized and rehydrated, sections were heated using an autoclave in 0.01 mol/L citrate buffer (pH 7.0) for antigen retrieval. Sections were incubated with 0.3% H2O2 in absolute methanol for 30 minutes to block endogenous peroxidase activity. Sections were then incubated with skimmed milk for 30 minutes to block non-specific staining. After this blocking step, sections were incubated with the primary antibodies shown in Table 2 at 4°C overnight. This was followed by sequential 1-hour incubations with the secondary antibodies (En Vision+ System-HRP-Labeled Polymer; Dako) and visualization with the Liquid DAB+ Substrate Chromogen System (Dako). All slides were counterstained with hematoxylin for 30 seconds before being dehydrated and mounted. We evaluated IHC results based on staining intensity and the percentage of positively stained tumor cells. The percentage of positively stained tumor cells was divided into five groups: no staining, <5% tumor cells reactive, 5–25% reactive, 25-50% reactive, and >50% reactive. The staining intensity level was divided into three groups: negative, weak, and strong. We designed a table to allocate IHC scores to each specimen. IHC scores were classified into three groups: negative (0), weak positive (1+), and positive (2+). We defined a 2+ IHC score as significantly positive. Scoring was simultaneously performed by two independent observers who were blinded to patient details.

### qRT-PCR

Total RNA was isolated using an RNeasy Mini Kit (Qiagen, Germany) in accordance with the manufacturer’s instructions. cDNA was produced using a ReverTra Ace qPCR RT-kit (Toyobo, Osaka, Japan), according to the manufacturer’s instructions. A list of the primers used is shown in Table 3. cDNA was subjected to quantitative SYBR Green real-time PCR by using SYBR Premix Ex Taq II (Takara Bio). A list of the specific primers used is shown in Table 3. qRT-PCR was performed with a LightCycler® Nano (Roche Diagnostic K.K.) using 40 cycles of a three-stage program with the following conditions: 20 seconds at 94°C, 40 seconds at 60°C, and 15 seconds at 72°C, as recommended by the manufacturer. The products were quantified during the initial exponential phase of amplification above the baseline. Data were obtained from triplicate reactions. The means and SDs of the copy numbers were normalized to the value for *glyceraldehyde-3-phosphate dehydrogenase* (*GAPDH*) mRNA.

**Table 3.**
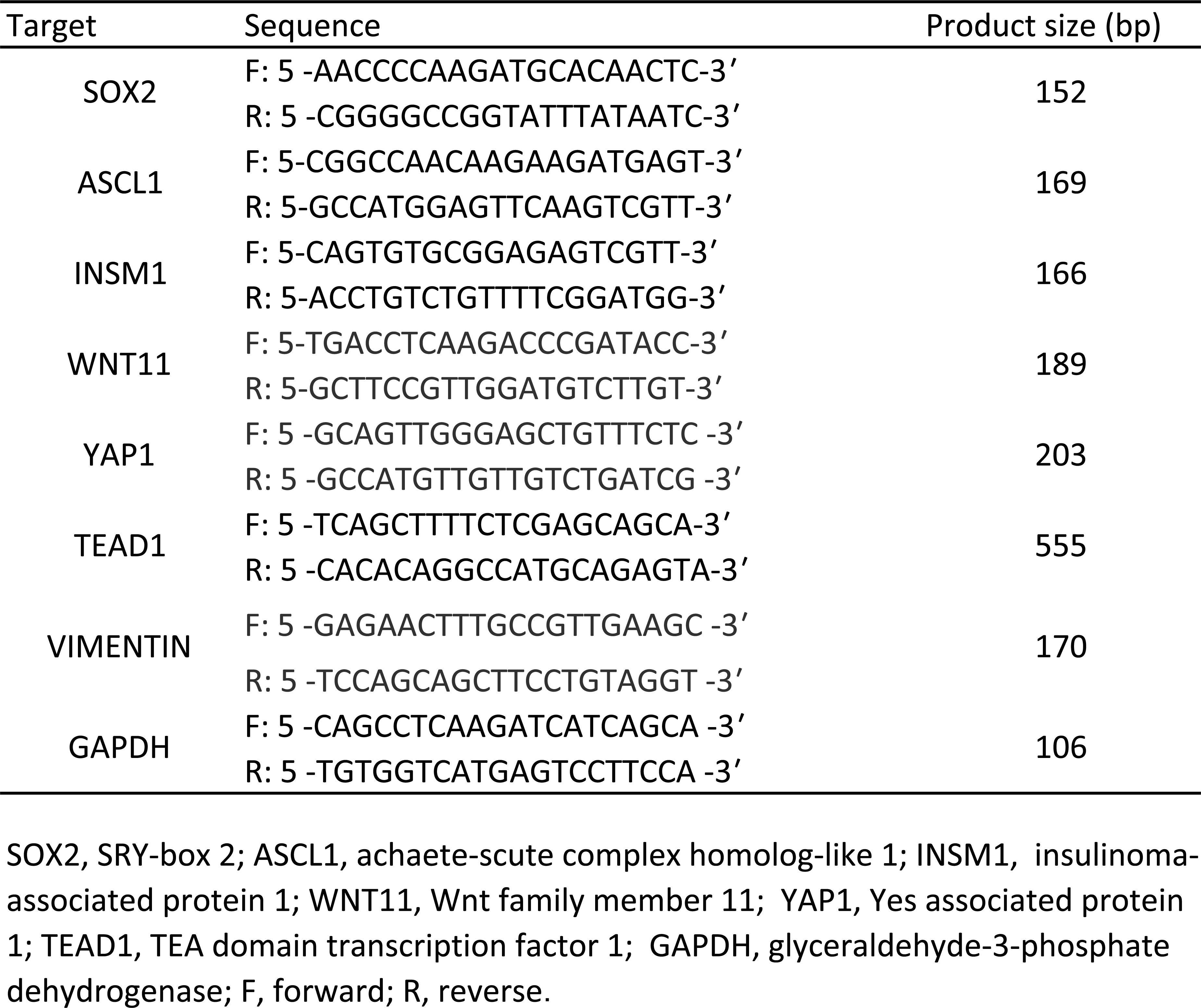
List of primers used in PCR

### *SOX2*-knockout experiment using SBC3 and H69AR cell lines

Genome editing using CRISPR/Cas9 was used for the knockout of the *SOX2* gene in the SBC3 and H69AR cell lines. We purchased a *SOX2*-knockout vector from GeneCopoeia^TM^ (Rockville, USA). These were 3 different human SOX2 sgRNA/Cas9 all-in-one expression clones (NM_003106.2). The sgRNA target sequences of *SOX2* were as follows: ATGGGCCGCTTGACGCGGTC (HCP217628-CG10-3-10-a), CGCCCGCATGTACAACATGA (HCP217628-CG10-3-10-b), and ATTATAAATACCGGCCCCGG (HCP217628-CG10-3-10-c). CCPCTR01-CG10 (GeneCopoeia^TM^) was used as a control. The sgRNA target sequence was as follows: GGCTTCGCGCCGTAGTCTTA. Regarding the establishment of *SOX2*-knockout H69AR cells, we obtained pSpCas9(BB)-2A-Puro(px459) from Addgene (Cambridge, MA) (Ran *et al.,* 2013). The sgRNA target sequences of *SOX2* were as follows: ATAATAACAATCATCGGCGG, GACCGCGTCAAGCGGCCCAT and ACAGCCCGGACCGCGTCAAG. Cells were harvested 48-72 hr post-transfection. These plasmids were co-transfected with Lipofectamine 3000 (Thermo Fisher Scientific) into cells at subconfluency. After 48 hr, transfected cells were treated with 200 µg/mL hygromycin B (Nacalai Tesque, Kyoto, Japan) or 1 µg/mL puromycin (Clontech) for the selection of stably transfected cells.

### Tumor xenotransplantation experiment

Eight-week-old male Rag2(−/−):Jak3(−/−) mice (a generous gift from Prof. S. Okada; Kumamoto University) were used. Two groups of mice were subcutaneously injected; one group was injected with 2 × 10^6^ stably transfected cells with *ASCL1*, and the other group was injected with an equal number of the control cell population. After 4 weeks, subcutaneous tumors were removed and fixed. The samples were fixed with 10% formalin and embedded in paraffin. Tissue sets were stained with hematoxylin and eosin and additional sections were used for IHC staining. Regarding tumor xenograft growth, a total of 1.0×10^6^ cells each of the mock-transfected and *SOX2* knockout SBC3 cell lines and mock cells were subcutaneously injected into the backs of mice. Twenty days after the first injection, tumors were removed and measured. All animal experiments were performed in accordance with the Institutional Animal Care and Use Committee guidelines.

### *In situ* precursor SCLC model mouse

We established a genetically engineered mouse model for *in situ* SCLC precursor lesions. These mice are *Trp53* gene-deficient, have Clara cell secretory protein promoter (CCSP) rt TA, tetO Cre-recombinase, floxed *Rb1*, and floxed *Hes1* genes, and were kept under the hypothesis that tumorigenesis of SCLC may occur with the double knockout of the suppressor oncogenes, *p53* and *Rb1*, and inactivation of the Notch signal pathway (Meder *et al.,* 2016). *P53* KO mice (ICR.Cg.-*Trp53*<TM1SIA>/Rbrc) (Tsukuda T *et al.,* 1993) were obtained from the Riken BioResource Center (Tsukuba, Japan), CCSPrtTA; tetOCre mice (Tichelarr *et al.,* 2002) were a generous gift from Dr. J. Whitsett. Floxed *Rb1* mice (FVB;129-Rb1^tm2Brn^/Nci; Vooijs *et al.,* 1999) were obtained from the NCI Mouse Repository (Frederick, MD), and floxed *Hes1* mice (Imayoshi *et al.,* 2008) from Dr. R. Kageyama of Kyoto University. Animals were kept under standard laboratory conditions with free access to water and food, and were maintained on a 12-h light/dark cycle under pathogen-free conditions. Doxycycline (Sigma-Aldrich) was dissolved in drinking water at a concentration of 1 g/L, and given to 4-week-old male animals for 6 weeks. After the treatment with doxycycline, animals were sacrificed with an intraperitoneal injection of an overdose of pentobarbital, and lung tissues were fixed in phosphate-buffered fixed 4% paraformaldehyde for 1 week. Fixed lung tissues were embedded in paraffin, and paraffin sections were used for hematoxylin eosin staining and immunostaining for Ascl1, Sox2, and Insm1. The present study was approved by the Animal Care Committee of Kumamoto University (#A2019-038).

### Statistical analysis

Spearman’s correlation coefficient (ρ) was calculated for the correlation analysis. Cell counting data were expressed as the means ± standard deviation of triplicate measurements. Differences in mean values between the groups were analyzed by a two-tailed statistical analysis using the Student’s *t*-test. GraphPad Prism version 7.04 (San Diego, CA) was used for the statistical analysis. *p* values less than 0.05 were considered to be significant.

### Dataset availability

The dataset produced in this study is available in the following database: ・ RNA-seq data: Gene Expression Omnibus GSE60052 (https://www.ncbi.nlm.nih.gov/geo/query/acc.cgi?acc=GSE60052)

## Acknowledgments

We thank Ms. Yuko Fukuchi and Ms. Takako Maeda for their technical assistance; the staff of LILA for their technical support; the Institute of Molecular Embryology and Genetics, Kumamoto University for its help with the RNA-seq analysis. This study was partially supported by a Grant-in-Aid for Scientific Research from the Ministry of Education, Culture, Sports, Science and Technology of Japan 18K19480 and by Aihara Pediatric and Allergy Clinic, Yokohama, Japan. This study was also supported in part by the program of the Joint Usage/Research Center for Developmental Medicine, Institute of Molecular Embryology and Genetics, Kumamoto University.

---

**Supplementary Fig. S1:**
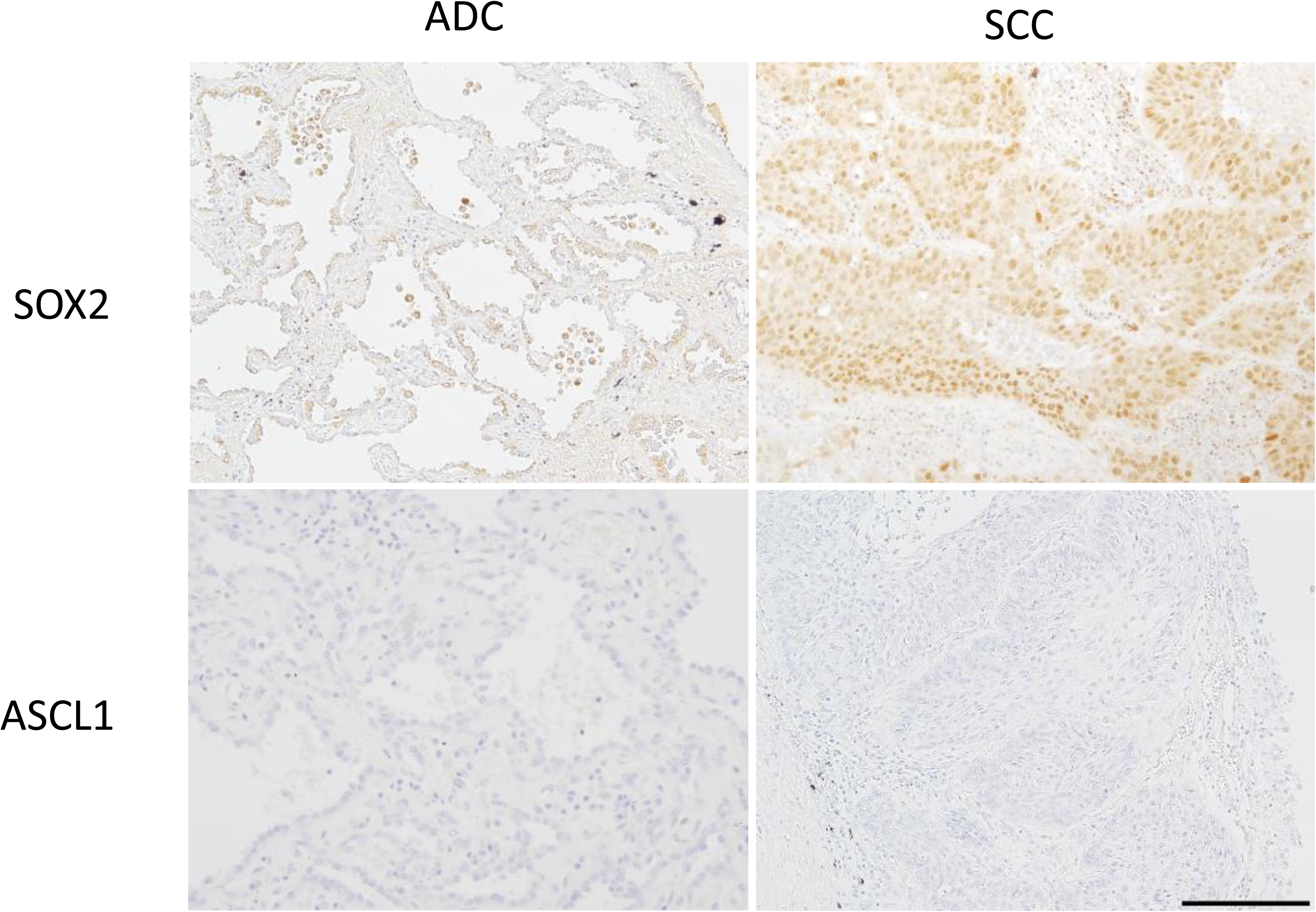
IHC images of surgically resected non-SCLC tissues for SOX2, and ASCL1. SOX2 was expressed in tumor cell nuclei. Representative images are shown. Scale bar = 200 μm.

**Supplementary Fig. S2:**
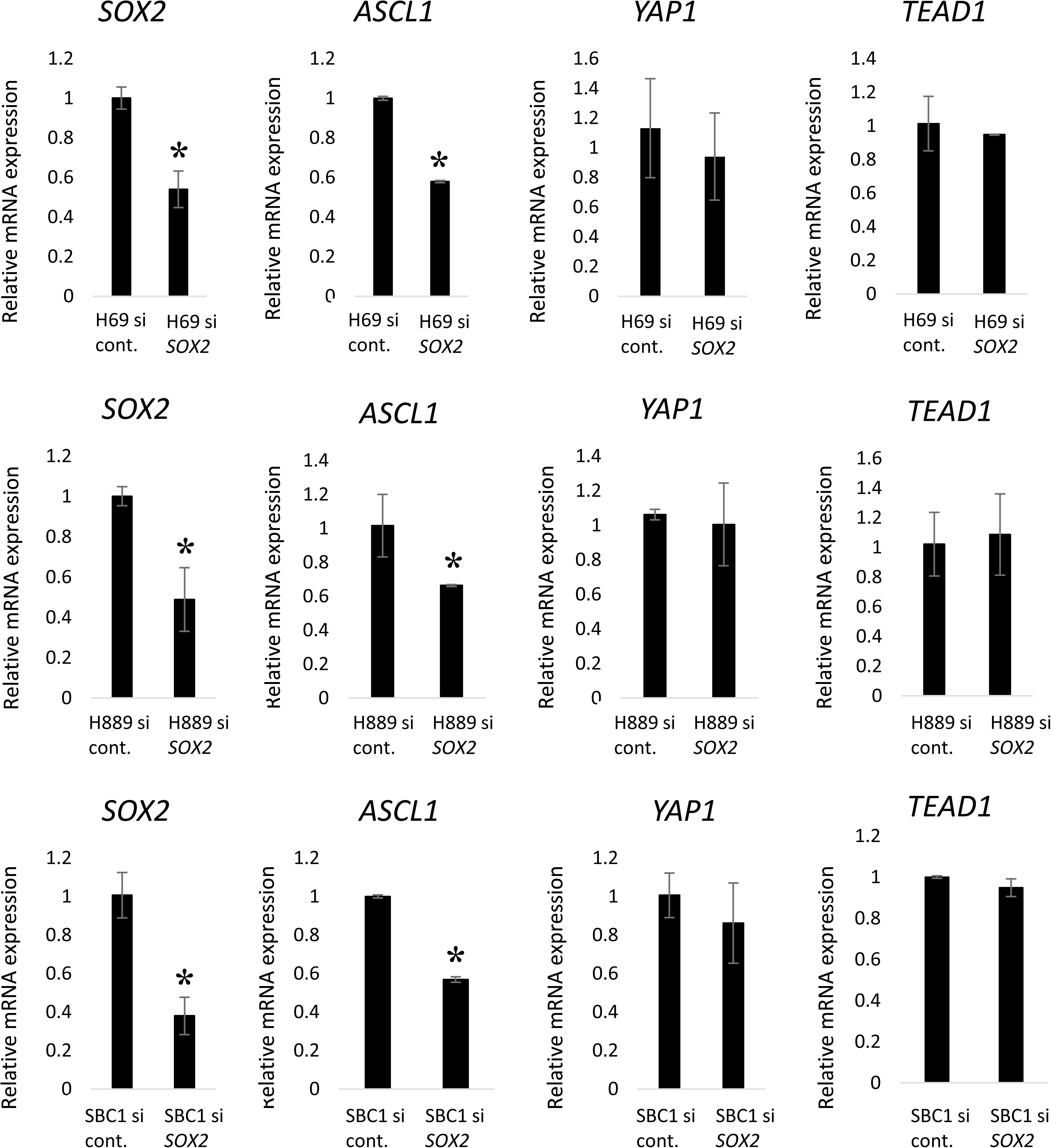
SOX2 affects ASCL1 expression in SCLC cell lines. The suppression of *ASCL1* was observed in the classical subtype of SCLC cell lines, H69, H889, and SBC1, with RNAi for *SOX2* by qRT-PCR. Hippo pathway-related mRNA, *YAP1* and *TEAD1*, did not show significant changes after the knockdown of *SOX2* in these cells. The analysis was performed in triplicate. Data are shown as the mean ±SD. Asterisks indicate a significant difference. *, p<0.05.

**Supplementary Fig. S3:**
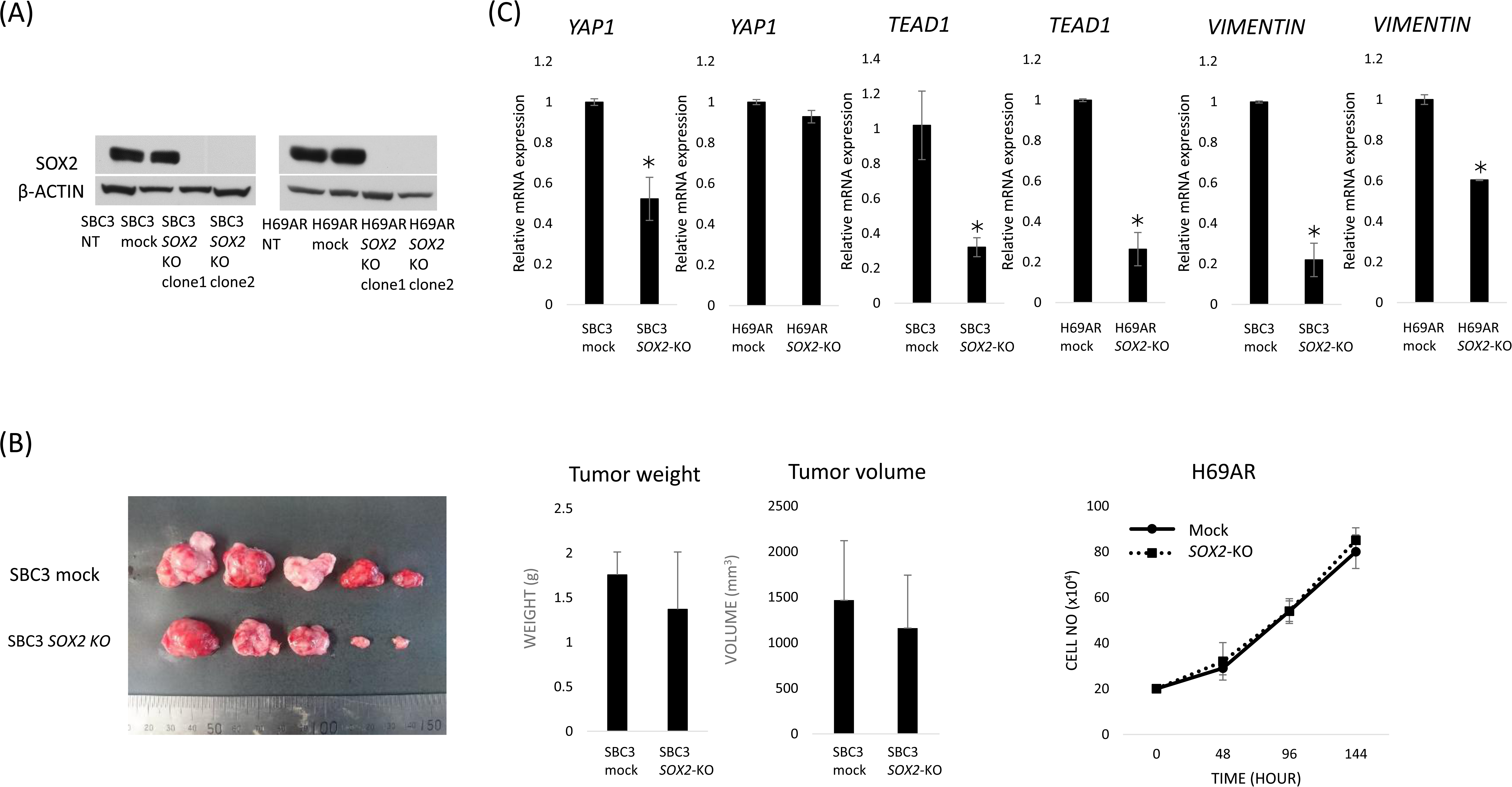
*SOX2* knockout in SBC3 and H69AR cells using the CRISPR/Cas9 system. SOX2 protein expression was completely diminished (A), whereas tumor proliferative capacity was not after the knockout of *SOX2* in SBC3 and H69AR cells (B). *YAP1*, *TEAD1,* or *VIMENTIN* mRNA expression decreased in SOX2-knockout SBC3 and H69AR cells (C).

## The paper explained

### PROBLEM

Small cell lung cancer (SCLC) is an aggressive neuroendocrine (NE) malignancy with few therapeutic options. *SOX2* is an oncogene, the amplification of which has been reported in human SCLC. However, the role of SOX2 remains unclear and strategies to selectively target SCLC cells have not been established.

### RESULTS

A chromatin immunoprecipitation sequencing analysis identified distinct putative target genes of SOX2 between the classical and variant subtypes of SCLC cell lines. ASCL1, a lineage-specific transcriptional factor, is involved in NE differentiation and tumorigenesis, and recruited SOX2 to the lineage-specific gene, *INSM1*, in the classical subtype of SCLC. *SOX2* suppression using RNAi resulted in significant reductions in tumor cell proliferative capacity in the classical subtype. On the other hand, SOX2 binds distinct genes, such as those in the Hippo signaling pathway, and the knockdown of *SOX2* was insufficient to inhibit tumor cell growth in the variant subtype of SCLC. SOX2 appears to potently modulate epithelial-to-mesenchymal transition via the Wnt or Hippo signaling pathways and affect tumor cell invasive capacity in each cell in a context-dependent manner.

### IMPACT

The present results provide insights into the functional underpinnings of small cell lung carcinogenesis and promote our understanding of SCLC phenotypic changes. The functional discrepancy in SOX2 between the classical and variant subtypes of SCLC may have an impact on appropriate therapeutic approaches and suggests that the ASCL1-SOX2 axis is a promising therapeutic target or biomarker to identify SCLC patients that may benefit from a SOX2-directed therapeutic approach in future clinical trials.

